# Perception of speech rate and intensity in Parkinson’s disease

**DOI:** 10.64898/2026.05.13.724886

**Authors:** Mishaela DiNino, Christopher C. Heffner, Kris Tjaden

**Affiliations:** Department of Communicative Disorders and Sciences, University at Buffalo, Buffalo, NY 14214; Center for Cognitive Science, University at Buffalo, Buffalo, NY 14260

**Author notes:** Correspondence should be addressed to Mishaela DiNino, 122 Cary Hall, South Campus, Buffalo, NY 14214.

**Keywords:** Parkinson’s disease, speech perception, learning

## Abstract

**Purpose:** Parkinson’s disease (PD) is a neurodegenerative disease that affects motor control but can also influence sensory perception. Changes in vision and proprioception are well-documented but less is known about how PD alters auditory perception, particularly perception of speech acoustic properties. The current study examined perception of speech rate and intensity in PD and the relationship of auditory perception to disease severity.

**Method:** People with PD were compared to age- and hearing-matched controls using perceptual tasks focused on discrimination and learning of speech rate and intensity. For rate discrimination, speech, non-speech, and visual stimuli were included to determine whether performance differences for PD participants and controls were specific to speech. Disease severity was assessed using the MDS-Unified Parkinson’s Disease Rating Scale (MDS-UPDRS) and the relationship to performance on perceptual discrimination and learning tasks was evaluated.

**Results:** People with PD performed significantly worse than controls in the rate discrimination task for all types of stimuli. There were no significant group differences for intensity discrimination. However, participants with greater PD disease severity demonstrated significantly poorer intensity discrimination accuracy. Performance on learning tasks utilizing rate and intensity manipulations did not differ between PD and control participants and was unrelated to PD disease severity.

**Conclusions:** People with PD had difficulty discriminating rate differences across speech, non-speech, and visual stimuli, indicating that challenges with rate perception are not limited to speech. The relationship between intensity discrimination and disease severity may suggest common dopaminergic networks between motor symptoms and auditory perception in PD.

## Introduction

Parkinson’s disease (PD) is one of the most common neurodegenerative diseases. Its prevalence is increasing largely due to international demographic shifts (Ben-Shlomo et al., 2024), necessitating a thorough understanding of its consequences. Cardinal motor symptoms of PD in the form of slowed movement speeds (bradykinesia), reduced movement amplitudes (hypokinesia), muscle stiffness (rigidity), postural instability and resting tremor are well documented (see Jankovic, 2008 for review). Yet, people with PD also experience non-motor symptoms. Many previous investigations have found differences in sensory perception between participants with PD and controls. Prior studies have also observed impaired category learning of sensory information in people with PD (e.g., Maddox et al., 2005; Shohamy et al., 2004). Given the centrality of effective communication to daily life, its role in social relationships, educational outcomes, occupational achievement, and participation in society, the investigation of learning and perception of speech acoustic characteristics in PD are of particular interest. The present study examined the effects of PD on learning to categorize sentences differing in rate and intensity and on discrimination of sentences based on those acoustic properties.

### Learning in Parkinson’s Disease

PD is characterized by dopaminergic neuron degeneration and subsequent abnormalities in the circuitry of the basal ganglia (see Blandini et al., 2000 for review), structures that have been linked to both learning and memory functions (see Packard and Knowlton, 2002 for review). Speech perception requires listeners to learn, adjusting their existing phonetic categories to detect speech variations such as dialect or rate, or to learn new categories in a second language. Dopaminergic systems in the basal ganglia may support these abilities (Heffner & Myers, 2021; Lim et al., 2014) but are impaired in Parkinson’s disease (Foerde & Shohamy, 2011; Shohamy et al., 2008).

Previous studies have suggested that difficulties with learning—especially procedural learning, a type of skill learning that happens with routine practice and is typically subconscious—are common in PD (Krebs et al., 2001). The learning of new motor routines is especially challenging (Marinelli et al., 2017). Perceptual category learning, which depends on procedural learning, is also impaired in PD. In visual procedural learning tasks, participants learn to categorize items based on the length of line segments or the presence of certain visual features. People with PD show difficulty relative to age-matched controls in their ability to learn visual categories (Ashby et al., 2003; Filoteo & Maddox, 2014; Maddox et al., 2005; Shohamy et al., 2004).

In the auditory domain, Heffner et al., (2022) found that people with PD performed significantly worse than controls adapting to rate-compressed speech. Perceptual adaptation involves adjusting to changes in auditory signals, thereby recruiting learning systems. Still, category learning of speech characteristics at the sentence level, the most ecologically relevant speech stimuli, had yet to be examined in people with PD.

### Sensory Perception in Parkinson’s Disease

PD is primarily a motor disorder, but at least some motor challenges experienced by individuals with PD may stem from sensory issues or impaired integration between sensory and motor information. Relative to healthy controls, people with PD experience greater difficulty completing tasks that require proprioception, defined as perception of where the body is in space (Jobst et al., 1997). Proprioception is necessary for accurate targeted muscle movements but declines with increased PD severity and disease duration (Maschke et al., 2003), suggesting that decreased proprioception contributes to motor dysfunction in PD (see Konczak et al., 2009 for review). In addition, people with PD can have vestibular dysfunction (e.g., Reichert et al., 1982; Rosado-Martins et al., 2025). Although evidence is mixed, some studies suggest that postural instability in PD arises from reduced integration between vestibular and visual perceptual systems in these individuals (see Smith, 2018 for review). The motor symptoms of PD may indeed stem from reduced ability to integrate sensory and proprioceptive information when planning and executing motor movements (Schneider et al., 1987; Almeida et al., 2005).

Altered sensory perception in PD, however, is not limited to systems associated with motor function. Some people with PD experience reduced odor perception due to changes in the olfactory system (Hawkes et al., 1997; for review see Fullard et al., 2017). People with PD have also been found to exhibit enhanced sensitivity to pain (Thompson et al., 2017) and reduced sensitivity to visual flickers and spatial contrast (Bodis-Wollner et al., 1987). Other research has shown that PD participants, compared to controls, demonstrate poorer temporal discrimination thresholds for somatosensory stimuli (see Lee et al., 2018 for review) due to the basal ganglia’s involvement in timing perception (see Yin et al., 2014 for review). Temporal discrimination impairment in people with PD has also been found in other sensory domains. In one investigation, people with PD required greater time differences between successively presented tactile, visual, and auditory stimuli to perceive two separate stimuli rather than one. Temporal discrimination thresholds correlated with disease severity and improved with medication, suggesting that changes in the dopaminergic system contributed to both sensory and motor impairment in PD (Artieda et al., 1992).

### Auditory Perception in Parkinson’s Disease

Several studies have demonstrated changes in auditory perception with PD, particularly in auditory perception of timing, rate, and rhythm. In one experiment, participants heard a repeated pair of rhythmic tones before hearing a third sequence that was either the same or different from the first pair. People with PD performed significantly worse than controls, particularly on simple rhythmic contrasts (Grahn & Brett, 2009). In another study of non-linguistic timing perception, participants with PD were asked to judge whether a tone was “short” or “long”. Participants who were less precise or less accurate generally had a PD diagnosis of at least 4 years old, a diagnosis of depression, and were taking a dopamine-enhancing medication (DiMarco et al., 2023). Participants in that investigation performed the task on two separate occasions: once after taking their dopamine-based medication as normal (“on medication”), and once after refraining from taking medication 8 hours prior to the appointment (“off medication”). Participants did not significantly differ in their performance between sessions, indicating that dopamine-based medications did not improve the behavioral deficits in non-linguistic timing perception. Whether individuals with PD also experience alterations in perceiving acoustic cues of speech timing remains unclear.

Accurate auditory perception is critical for successful communication, but prior studies have found impairments in perception of the acoustic properties of speech in people with PD relative to healthy controls. Previous research on speech perception in PD has generally focused on three attributes of speech: emotional prosody, intensity, and speech rate (Kwan & Whitehill, 2011), which all require perception of specific acoustic properties of the auditory signal. As discussed below, these studies have often found differences between people with and without PD. It is worth noting that differences in performance on auditory tasks cannot solely be attributed to sensorineural hearing loss in the PD population, a topic of recent research (da Silva Lopes et al., 2018; Liu et al., 2020; Jose et al., 2024; DiNino et al., 2026).

Emotional prosody, defined as the patterns of rate, intensity, fundamental frequency, and voice quality that signal the emotional state of the talker, has been widely studied in PD. People with PD struggle to differentiate emotions based on prosodic cues alone (Schröder et al., 2010). This is especially true for negative emotions (Dara et al., 2008). Studies of cortical activity in people with PD further indicate that sad prosody is processed differently in people with PD than controls (Dara et al., 2008; Schröder et al., 2006). The multifaceted nature of emotional prosody, however, poses challenges for research. It is only through a combination of acoustic cues that listeners extract the emotional state of their interlocutors, which means that studies of emotional prosody cannot isolate the influence of individual acoustic cues without utilizing advanced digital signal processing techniques.

Other studies have focused on perception of individual acoustic cues, such as intensity. Compared to healthy controls, people with PD perceive the loudness of speech produced by others (i.e., extraphonic speech) as well as loudness of their own speech (i.e., autophonic speech) differently. For example, individuals with PD tend to perceive objectively quiet speech as louder than healthy controls, especially for autophonic speech (Clark et al., 2014; De Keyser et al., 2016; Ho et al., 2000). People with PD have also been found to differ from healthy controls in the ability to distinguish speech sounds differing in intensity. For example, Richardson and Sussman (2019) assessed people with PD and age-matched controls on a set of discrimination tasks containing a steady-state [u] vowel. People with PD performed significantly worse than age-matched controls on the discrimination tasks for a wide range of intensity differences, with the largest differences generally present for stimuli differing by 4 dB SPL.

Speech rate refers to the pace at which an individual speaks, including pauses, and is an acoustic property of speech that has been much less commonly studied in investigations of perceptual abilities in PD. This speech characteristic is of particular interest in PD because the basal ganglia, which have impaired neuronal circuitry in PD, play a role in timing perception (see Yin et al., 2014 for review). The only relevant study to date examined perception of monosyllabic words in sentences that were spoken with either conversational, slow, or fast rates in participants with PD and neurologically healthy controls of similar ages (Forrest et al., 1998). Results indicated that people with PD performed significantly worse than controls on word recognition at the slow speech rate. However, on average, PD participants also had greater difficulty identifying the words at conversational and fast rates relative to controls, suggesting a general impairment in word recognition among those participants. In that study, talkers producing the rate-varying stimuli were asked to speak at particular rates; the current study provides stricter control over speech rate with physical speech stimulus manipulations to investigate speech rate perception in PD.

### Study Purpose

The current study sought to extend understanding of speech intensity and speech rate learning and perception in PD, processes that are both important for successful verbal communication. This line of inquiry also could help to shed light on the neurobiological underpinnings of speech perception, especially related to learning. The neurobiological hallmark of PD is basal ganglia dysfunction, which impairs procedural learning in people with PD (Shohamy et al., 2004). Studies have proposed that procedural learning in the basal ganglia is a prime contributor to speech perception (Heffner & Myers, 2021; Lim et al., 2014). Studying speech perception in PD, then, is a plausible avenue to evaluate those proposals. Finally, examining domain-specificity of timing issues could help determine whether these potential difficulties are unique to speech. Building upon previous research, the current study used both speech and non-speech stimuli to help determine the extent that timing and learning issues are shared across sensory domains.

The present study sought to advance understanding of auditory perception in PD by evaluating perception of acoustic properties of speech – arguably the most important auditory stimulus for everyday spoken communication. Specifically, we compared learning and discrimination of speech rate and intensity between people with PD and age- and hearing-matched controls. Based on the prior research described above, we hypothesized that people with PD would require more trials to learn categories and exhibit poorer discrimination of acoustic speech characteristics, and that these deficits should be present for both rate and intensity. We also assessed motor and non-motor symptoms in PD participants to examine how disease severity may relate to perception of speech rate and intensity.

## Methods

### Participants

A total of 71 participants (32 in the PD group, 39 without PD) were recruited through a university database and via word-of-mouth from other study participants. All participants were required to be between the ages of 45 and 80, native speakers of North American English, and no known neurological conditions other than PD. Use of a hearing aid or cochlear implant also excluded individuals from participating. PD participants were additionally required to have a PD diagnosis from a neurologist, as listed in the database that was used for recruitment. History of speech disorders was exclusionary for both groups. Control participants were recruited to match age, gender, and audiometric characteristics of individual PD participants. To identify potential participant matches, propensity score matching was performed using the MatchIt package (Ho et al., 2011) in R software (R Core Team, 2024). A distance score was calculated for each potential control-PD match using a combination of age, gender, and the audiometric pure-tone average (PTA; using hearing thresholds at 500, 1000, and 2000 Hz, see below). Matching sought to minimize the average distance score for people with PD and controls using an optimal pair matching algorithm (Hansen & Olsen Klopfer, 2006). Participants provided written informed consent at the beginning of the first study appointment. All study procedures were approved by the University at Buffalo Institutional Review Board (STUDY00005259).

One participant with PD reported that their neurologist was considering an alternative diagnosis and was excluded, and one control participant reporting a history of speech therapy and was excluded from the final sample. Additionally, one participant with PD and three controls were excluded for failing to return for a required second testing session. All participants were screened for mild cognitive impairment using the Montreal Cognitive Assessment (MoCA; Nasreddine et al., 2005). Participants with scores lower than 21 were excluded, following recent guidance for more permissive scoring than the standard score of 26 for a diagnosis of mild cognitive impairment (Dautzenberg et al., 2022). One participant with PD with a MoCA score below 21 was excluded. Six control participants were additionally excluded because other controls more closely matched PD subjects. The final sample consisted of 29 people with PD and 29 matched controls. Summary demographic information is presented in Table 1. It should be noted that the sample size is larger than many prior related studies, which ranged from 10 to 27 PD participants (Clark et al., 2014; De Keyser et al., 2016; Forrest et al., 1998; Heffner et al., 2022; Ho et al., 2000; Richardson & Sussman, 2019:19; Schröder et al., 2006).

**Table 1.** Demographic and speech characteristics of people with Parkinson’s disease (PD) and controls. Numbers in parentheses are standard deviations. PTA: pure tone average, calculated by averaging thresholds across ears for 500, 1000, and 2000 Hz. dB: decibels. HL: hearing level. MoCA: Montreal Cognitive Assessment.

|  | People with PD | Controls |
| --- | --- | --- |
| Gender | 11 Women<br>18 Men | 15 Women<br>14 Men |
| Race | 23 White<br>1 Black<br>5 Not Reported | 27 White<br>1 Black<br>1 Not Reported |
| Age (years) | 71.0 (6.55) | 70.1 (5.62) |
| PTAs (dB HL) | +15.1 (10.3) | +13.4 (8.36) |
| Years of Formal Education (years) | 15.9 (2.75) | 16.0 (2.49) |
| MoCA Score | 25.3 (2.69) | 25.7 (2.10) |
| Speech Severity | 10.6 (7.11) | 6.2 (4.40) |
| Speech Intelligibility | 97.8 (2.36) | 99.4 (0.62) |
| Speech Listening Effort | 5.8 (4.71) | 2.7 (1.81) |

Ratings of speech severity, intelligibility, and listening effort for the first six sentences of the Rainbow Passage (Fairbanks, 1960) were obtained for all speakers to document the presence of perceptible speech impairment. Participants were audio recorded reading aloud the Rainbow Passage. Passages for seven participants with PD and two controls were not recorded due to technical errors or were excluded due to significant distortion. For the remaining passages, three speech-language pathologists (SLPs) rated intelligibility (“rate the understandability of the speech”), listener effort (“how effortful was it for you to understand the speaker?”), and speech severity (“rate the severity of the speech”) on a computerized visual analogue scale (VAS) ranging from 0 to 100, with higher values reflecting greater intelligibility, increased listening effort, and greater speech severity. Reading passages were randomized and the SLPs were blinded to speaker group identity. The order in which SLPs judged intelligibility, listener effort and severity was counterbalanced, with intelligibility always preceding listener effort. The order in which a given SLP provided VAS ratings of intelligibility, listener effort and severity remained the same for all passages. Summary data are reported in Table 1.

Shapiro-Wilk normality tests indicated that the VAS rating data were not normally distributed and therefore Wilcoxon rank sum tests were performed to determine any group differences. The PD group was rated as more severe (*W* = 430.5, *p* = .008), less intelligible (*W* = 134.5, *p* = .001), and required more effort to understand (*W =* 429, *p* = .007). While statistically significant, these group differences are likely not clinically meaningful (see Stipancic et al., 2025). The very mild nature of any speech impairment or dysarthria in the PD group is considered further in the discussion. Table 1 includes participant demographic information as well as group means and standard deviations for VAS ratings of intelligibility, listener effort, and speech severity.

### Audiometric Threshold Testing

Pure tone audiometry was conducted in a sound-attenuating booth using the Hughson-Westlake “up-5, down-10” procedure. Participants heard tones from 250-8000 Hz in octaves through RadioEar IP30 insert earphones and responded by pressing a button when they detected a tone. Threshold was determined at each frequency to be the intensity (in dB HL) at which the participant responded on at least two out of three presentations. Hearing thresholds at extended high frequencies (10k-16k Hz) were also collected and are reported in DiNino et al. (2026). It should be noted that the PD and control groups were statistically similar in their extended high frequency hearing thresholds (see DiNino et al., 2026), and therefore no group threshold differences at those frequencies would influence group-level performance on the speech rate and intensity tasks.

The pure tone average (PTA) was calculated for each participant by averaging thresholds at 500, 1000, and 2000 Hz across ears. PTAs were then used to match controls with PD participants. Average pure tone hearing thresholds for each group are shown in Figure 1.

**Figure 1.**
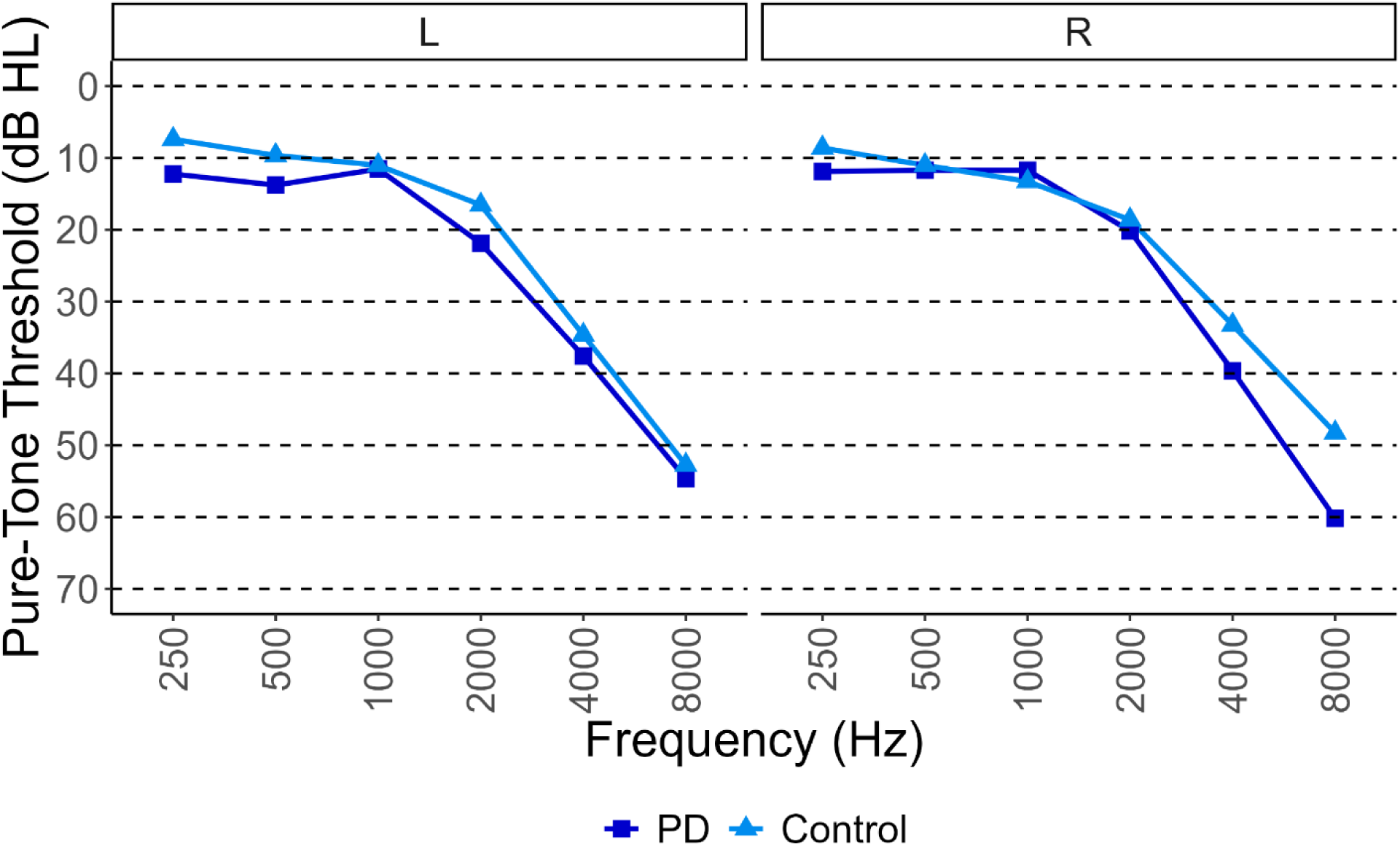
Average hearing thresholds between 250 Hz and 8000 Hz for the people with Parkinson’s disease (PD; dark blue squares) and controls (light blue triangles), split by left (L) and right (R) ear.

### Assessment of Disease Severity

Disease severity for participants with PD was assessed using the Movement Disorder Society – Unified Parkinson’s Disease Rating Scale (MDS-UPDRS) (Goetz et al., 2008). In Part I, participants report experiences with thirteen non-motor symptoms such as cognitive impairment, hallucinations, mood changes, and issues with sleep. In Part II, participants report experiences with motor symptoms that may affect thirteen everyday activities such as speaking, eating, and getting dressed. Each question asks participants to rate whether they experience no problems in that domain (a score of 0) ranging to severe problems (a score of 4) in which PD symptoms in that domain cause problems all the time. Therefore, Parts I and II rely on participant report. In contrast, Part III is an experimenter assessment of motor function. The participant is asked to perform motor tasks such as speaking or moving their hand and the experimenter assigns a ratings ranging from 0 (“normal”) to 4 (“severe”). Part IV assesses involuntary muscle movements and “on” and “off” periods, which not all PD patients experience. Part IV was collected but not reported, as is common in studies utilizing the MDS-UPDRS.

Experimenters completed online MDS-UPDRS training prior to administration. Student research assistants collecting data additionally underwent in-person training and supervision led by C.C.H. or trained lab managers before they conducted the MDS-UPDRS independently.

### Stimuli for Learning and Discrimination Tasks

All materials used in the intensity and rate learning and discrimination tasks originated from a recording of the sentence “the puppies chased the ball,” which was produced on a single breath and contained no pauses or silent periods other than those associated with acoustic properties of the stimuli (e.g., closure interval for /p/ and /b/). The sentence was then modified using Praat (Boersma & Weenink, 2001) to create stimuli varying in intensity. The sentence was recorded by a cisgender female native speaker of North American English in a quiet, sound-attenuated room. The sound file was filtered to remove low-amplitude background noise via Praat’s “reduce noise” function. The sound was filtered from 80 Hz to 10kHz using spectral subtraction with a smoothing bandwidth of 40 Hz. Peak intensity was then rescaled to 70 dB SPL. From this base sentence, a variety of sentence versions were created that differed in peak intensity or duration (to alter perceived speaking rate). To create stimuli differing in intensity, Praat was used to scale the peak intensity to range between 62 and 78 dB SPL in 2-dB increments. To create items that differed in total duration and, by extension, speech rate, Pitch Synchronous Overlap and Add (PSOLA) was used in Praat to uniformly vary the duration of the sentence between 60% (fast) and 140% (slow) of the original sentence duration in 10% increments to create a continuum.

The learning tasks and the intensity discrimination task utilized the stimuli described above. The rate discrimination task included four types of stimuli (i.e., the rate-varying base sentences, reversed speech, non-linguistic tones, and visualizations) to determine whether any changes related to discrimination of stimulus rate were specific to speech or generalized to other domains. To create reversed speech, the acoustic content of the sentences was time-reversed in Praat to reverse the entire audio file. These items therefore contained all the timing and spectral information of the sentences, but lacked the lexical information found in the original recordings. To create non-linguistic tones, TextGrids in Praat were used to annotate the beginning and the end of the vowels in the filtered, original sentence. The vowels were replaced by a non-linguistic pure tone of 332 Hz (the median fundamental frequency of the original sentence) and the rest of the sentence was replaced with silence. Therefore, these stimuli consisted only of pure tones that broadly matched the duration of important timing-related components of the original sentence (e.g., the vowels) without containing speech information. Figure 2 shows the waveforms for each type of auditory stimulus.

**Figure 2.**
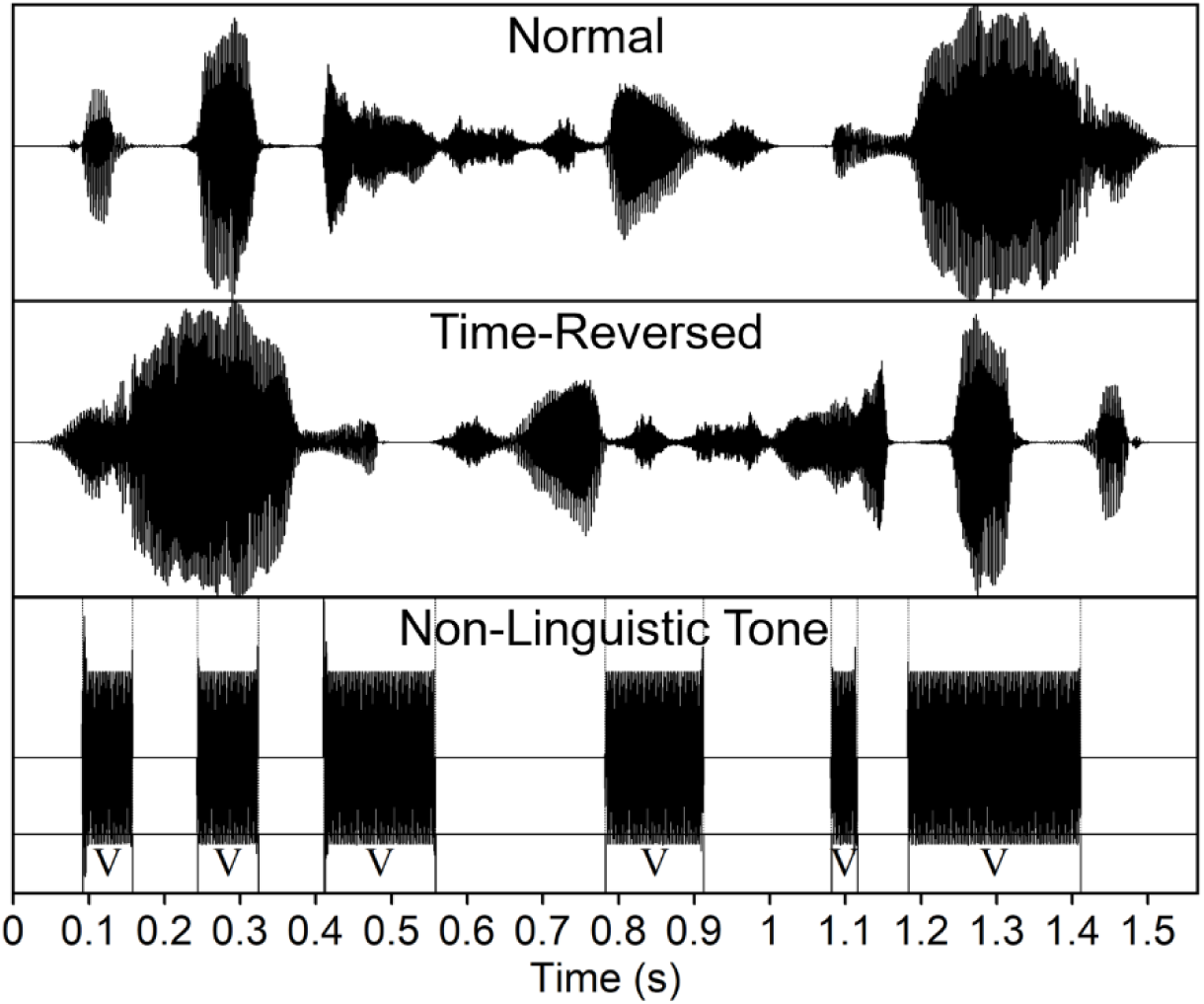
Waveforms of the auditory stimuli used in the rate discrimination task: normal speech (top), time-reversed speech (middle), and non-linguistic tones (bottom). The Praat TextGrid used to delineate the vowel portions of the original sentence is overlaid on the non-linguistic tone waveform. V: vowel portion of the stimulus.

To create the visual stimuli, the Headliner app (*Headliner App*, 2022) was used to generate a visualization of the base sentences. The app generated a time-varying spectrum of the original audio with a simple white-on-black color scheme (see Figure 3). Audio was removed from the video clips, meaning that participants could only discriminate stimuli based on the timing information in the visual stimuli.

**Figure 3.**
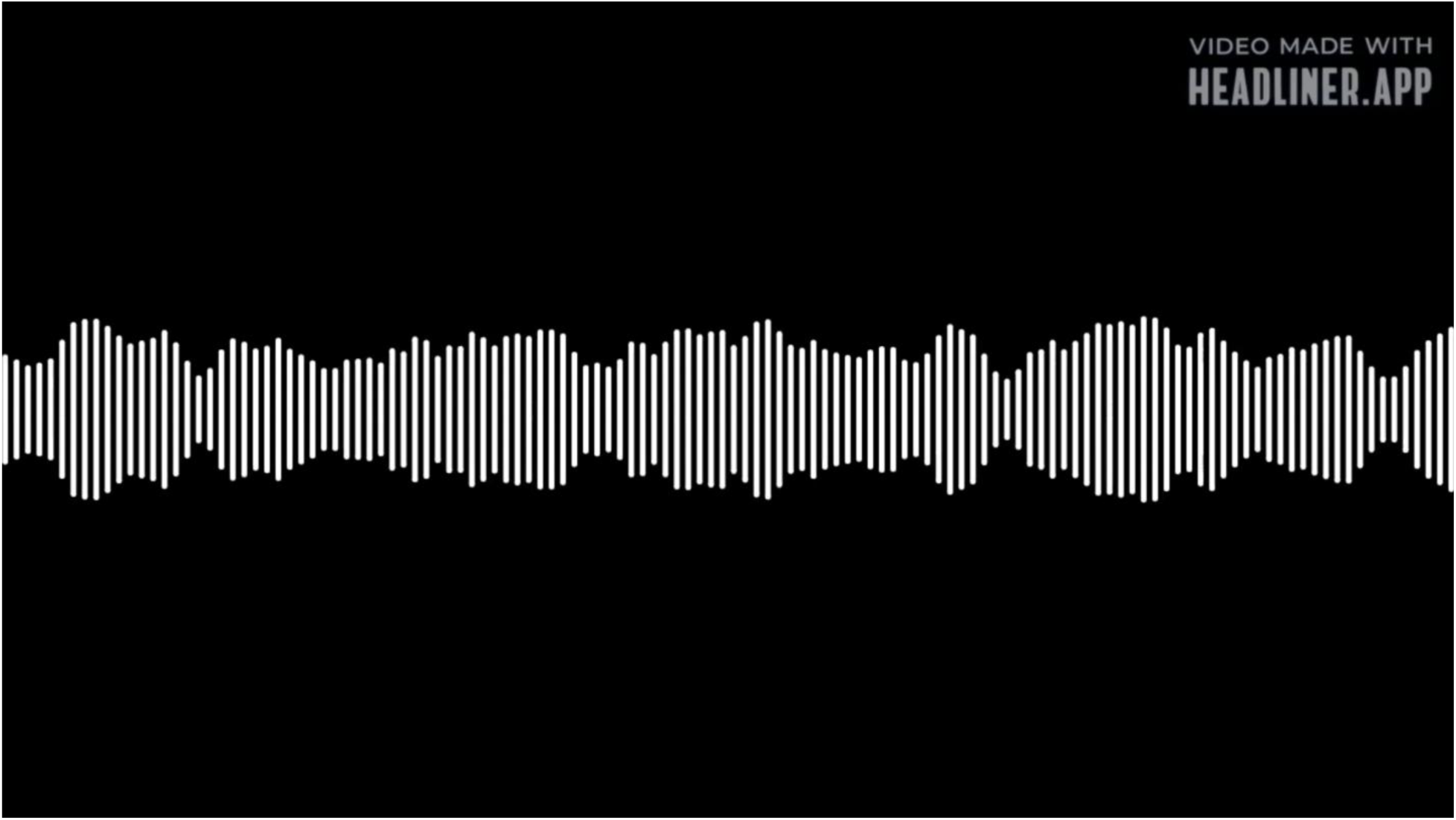
Screenshot of the stimuli for the visualization condition of the rate discrimination task. The spectrum (amplitude over frequency) shown is a visual representation of the auditory stimuli used in the task, created via the Headliner App. The visual spectrum changes over time throughout the duration of the sentence.

### Procedure for Learning Tasks

On each trial, participants heard one stimulus while viewing three squares of different colors (blue, yellow, and red) on the computer screen. After the sound was presented, participants used a button box to indicate which color square the sound “matched” with. They then received feedback about their selections: a check mark indicating a correct response or an X indicating an incorrect response. Participants were not told that the assignment of a stimulus to a particular color square was based on a particular speech rate or speech intensity. Instead, participants were instructed to use the feedback they received to guide their responses over the course of the experiment and learn to group the sentences. The learning task therefore relied on both the successful ability to distinguish sentences based on their rate and intensity as well as to learn from the feedback to use those distinctions productively. This task was modeled after successful previous studies of phonetic category learning (Heffner et al., 2019; Heffner & Myers, 2021).

Assignment of rate- or intensity manipulated stimuli to categories (“blue,” “yellow,” or “red” squares) followed similar structures across each of the two manipulated acoustic parameters. In the intensity learning task, items were separated by 2-dB intervals ranging from 62 to 78 dB. One category contained the lower intensity sounds (62, 64, and 66 dB), a second category contained the mid-intensity sounds (68, 70, and 72 dB) and a third category contained the higher-intensity sounds (74, 76, and 78 dB). In the rate learning task, items were separated by 10% intervals ranging from 60% of the original duration to 140% of the original duration. One category contained the faster sounds (60%, 70%, and 80% duration), a second category contained the approximately normal rate sounds (90%, 100%, and 110% duration) and a third category contained the slower sounds (120%, 130%, and 140% duration). Assignment of categories to square colors (i.e., whether any category was “blue”, “yellow”, or “red”) was assigned randomly for each participant. On a given trial, participants heard one sound and matched the sound with its assigned category, using the visual feedback to learn which category each particular sound was assigned to.

Participants completed one learning task containing only rate-manipulated stimuli and a second learning task containing only intensity manipulated stimuli. The order in which participants performed the learning tasks was counterbalanced. Each learning task (rate or intensity learning) consisted of 216 trials and took participants approximately 25 minutes to complete. Each task ended at the conclusion of the last trial.

### Procedure for Discrimination Tasks

Participants heard (or saw) versions of the stimulus played three times. The first (“A”) and third (“B”) instance of the target were always different. The second instance (“X”) matched either the first or third instance. Participants were asked to indicate whether X matched A or B by pressing a button on a response box. Correct responses indicated that the participant was able to use the acoustic cue being manipulated to distinguish between targets. Participants completed a four-trial practice block with non-linguistic stimuli that varied extensively in rate and intensity to familiarize themselves with the structure of the task before beginning the experimental trials.

For the intensity-based targets, the intensity of A and B always diverged by 4 dB. This value was based on Richardson and Sussman (2019), who found that a 4 dB difference in intensity was the point at which discrimination by people with PD and by age-matched controls was most dissimilar. The paired intensities were 62 and 66 dB, 66 and 70 dB, 70 and 74 dB, and 74 and 80 dB. Items were counterbalanced in terms of which target had a higher intensity than the other and whether X matched A or B. Ultimately, there were a total of 16 combinations of A, X, and B across the 16-dB range of intensities. The set of randomly ordered combinations was repeated 5 times for a total of 80 trials.

For the rate discrimination task, four types of stimuli (natural-speech, time-reversed, non-linguistic tones, and visual) were presented. Pilot testing was conducted for the rate discrimination task to find the difference in rate between A and B that matched the discriminability of the 4 dB difference in stimuli used for the intensity discrimination task. This difference was 20% duration, and therefore the rate of A and B always diverged by this amount on a given rate discrimination task trial. The paired rates were 60% and 80%, 80% and 100% (normal speech rate), 100% and 120%, and 120% and 140% duration. Again, items were counterbalanced in terms of the relative rate of A and B and whether X matched A or B. There was a total of 16 combinations of A, X, and B, repeated 5 times for a total of 80 trials for each stimulus type. The rate discrimination task contained one block for each stimulus type, totaling four blocks for which presentation order was randomized. Block order randomization was counterbalanced across participants.

Participants completed two discrimination tasks: one containing only rate-manipulated stimuli and one containing only intensity manipulated stimuli. The order in which participants performed the discrimination tasks was counterbalanced. The intensity discrimination task took participants 15-20 minutes to complete. The rate discrimination task, due to the four types of presented stimuli, took participants about 50 minutes to complete.

### Study Procedure

Tasks were completed across two experimental sessions each lasting 60-90 minutes, with the exception of one participant who required three sessions. Participants with PD were scheduled for sessions that began between 90 and 150 minutes after their most recent regularly scheduled dose of PD medication. In the first session, participants completed the MoCA followed by the MDS-UPDRS for participants with PD. Participants then completed audiometric threshold testing and categorical loudness scaling, a measure of how physical intensity maps to perceptual loudness among individuals. Detailed categorical loudness scaling methods and results are reported in DiNino et al. (2026). For three participants, audiometric threshold testing was performed in the second session due to unavailability of the sound-attenuating booth during their first session. All participants completed the learning and discrimination tasks in the second session. Learning tasks were administered before the discrimination tasks so that participants were not biased to the acoustic properties manipulated in the learning tasks.

Some participants had incomplete datasets, although the data that was collected is still reported here. One participant with PD withdrew due to fatigue during the discrimination portion of the experiment in Session 2 but completed the learning tasks. Experimenter error led to no data collected from the visual condition of the rate discrimination task from five participants (one with PD, four controls) and no learning data from one control participant. Most participants completed the two sessions within two weeks. Two participants who were initially excluded due to not meeting an initial, more restrictive MoCA score criterion were later included, which led to a lag of a few months between the first and second sessions.

### Analysis

#### Learning Tasks

All statistical analyses were performed using R statistical software (version 4.3.1). One analysis was conducted to test each study hypothesis. For each learning task, a mixed-model regression was conducted to predict trial accuracy over the course of the task. Participant group (control vs. PD), trial number (rescaled to range from 0 to 1 to aid in model convergence), and the interaction of group and trial number were included as fixed factors in each model to predict accuracy on each trial. The fixed factor of trial number indicates whether participants improved over time (i.e., whether they were learning); the fixed factor of group indicates whether there was an across-the-board difference between participant groups in accuracy; and the interaction term indicates whether either group learned faster over the course of the experiment. The fixed factors were complemented by a maximal set of random effects (Barr et al., 2013). This included random intercepts by participant, to allow participants to vary in their accuracy randomly, and by item, to allow some items to be easier to categorize than others (such as items far from category boundaries). Additionally, random slopes for trial number were included by participant to allow participants to vary randomly in their speed of learning across the experiment. Both regression models demonstrated convergence and nonsingular fits.

#### Discrimination Tasks

For the discrimination tasks, the dependent measure was *d’*, the sensitivity index commonly used in psychophysics (Hautus et al., 2021). In an AXB task, *d’* is calculated based on the A and X stimuli. If A and X are different and participants correctly indicate that they are different by responding “B” (meaning, X matches B), the response is a “hit,” as the participants are successfully able to discriminate between the sounds based on the acoustic parameter that was manipulated. If A and X are the same, but participants incorrectly indicate that they are different, the response is a “false alarm”, as the participants could not successfully use the acoustic parameter to distinguish the sounds that they heard. The participant’s response in this case would also be “B”, but in this case, the response would be incorrect. To obtain the sensitivity index, the hit rate and false alarm rate were converted to *z* scores, and the false alarm rate was subtracted from the hit rate. *d’* allows participants’ sensitivity to differences in rate or intensity to be separated from a bias towards responding “A” or “B” in this design. However, calculating *d’* requires pooling across all responses over the course of the experiment.

An independent-samples *t*-test was employed for the intensity discrimination measure to compare the PD and control participant groups, as only one stimulus type was utilized (e.g., sentences). A linear mixed-effect model was conducted for rate discrimination because the task included four stimulus types. Participant group (control vs. PD), stimulus type (normal speech, reversed speech, non-linguistic tones, and visualizations) and the interaction of group and stimulus type were included as fixed factors to predict *d’* for each participant. The model also included a random effect for participant to account for individual participant responses to the four different types of stimuli. Normal speech was set as the default speech stimulus type and the PD group was set as the default participant group.

#### Disease Severity and Task Performance

Linear modelling was conducted to assess the extent to which variability in PD symptom severity, as assessed by the MDS-UPDRS, explained variability in 1) intensity learning, 2) rate learning, 3) intensity discrimination, and 4) rate discrimination. Following Goetz et al. (2024) and DiNino et al. (2026), we used a composite measure that combined MDS-UPDRS Parts I (“non-motor experiences of daily living”) and II (“motor experiences of daily living”) as well as MDS-UPDRS Part III scores as independent variables in the models. Participants’ PTA, representing hearing thresholds at the frequencies important for speech perception (500, 1000, and 2000 Hz), were incorporated into each model as a covariate.

## Results

### Learning Tasks

#### Intensity Learning

Figure 4(A) shows average accuracy across the experiment in the intensity learning task. The generalized linear mixed model revealed a significant effect of trial number on accuracy (*β* = 1.09, *p* < 0.001). No significant effect of group (*β* = 0.14, *p* = 0.525) nor a significant interaction between trial number and group (*β* = 0.45, *p* = 0.206) were found.

**Figure 4.**
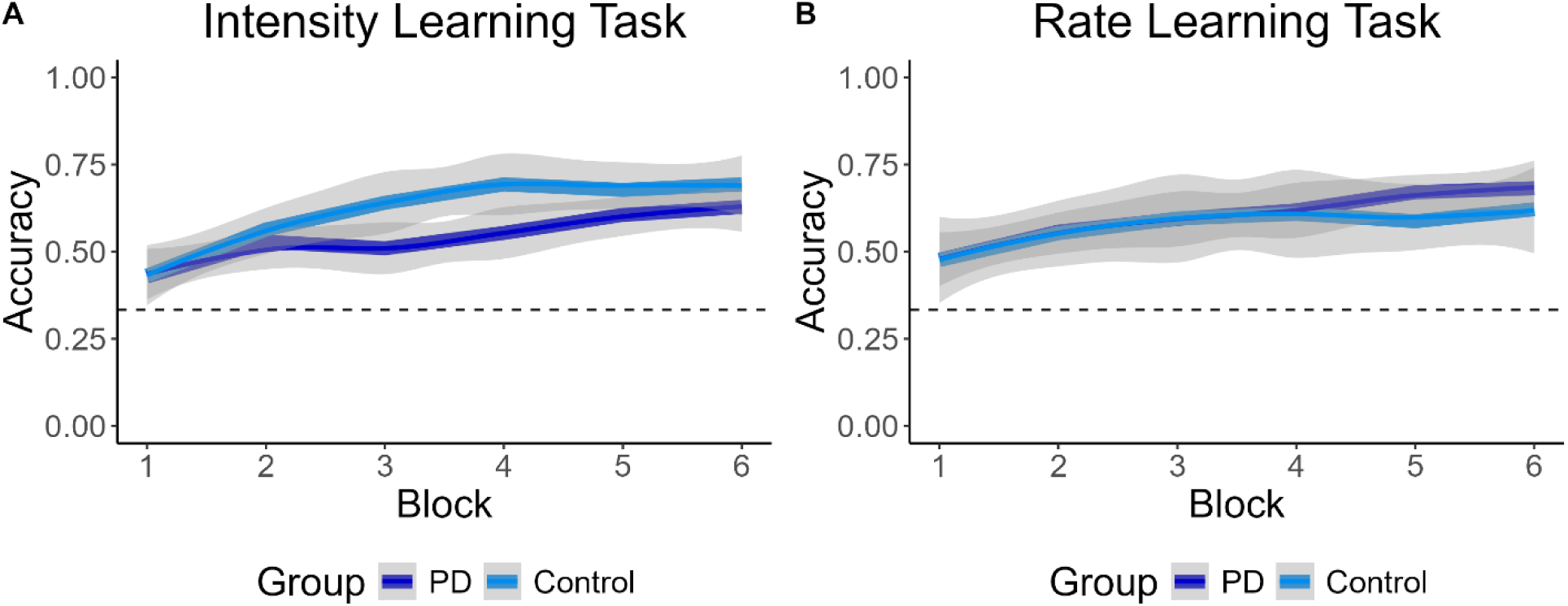
Average accuracy for each group (Participants with Parkinson’s disease [PD] in dark blue, controls in light blue) in the (A) intensity learning and (B) rate learning task. Gray shaded areas around each line indicate the 95% confidence interval of the mean for each group. Blocks indicate a set of one-sixth of the overall number of trials. The dashed horizontal line shows chance (33%).

#### Rate Learning

Figure 4(B) shows average accuracy across the experiment in the rate learning task. As with intensity learning, we observed a significant effect of trial number on accuracy (*β* = 1.33, *p* < 0.001), but no significant effect of participant group (*β* = 0.20, *p* = 0.578) or interaction between trial number and group (*β* = -0.52, *p* = 0.210).

### Discrimination Tasks

#### Intensity Discrimination

Figure 5 shows *d’* values for all participants. Although visual inspection of the data suggests a trend for participants with PD to have lower *d’* values for intensity discrimination relative to controls, an independent samples *t*-test indicated no significant difference for the two groups (*t*(49.7) = -1.78, *p* = .08).

**Figure 5.**
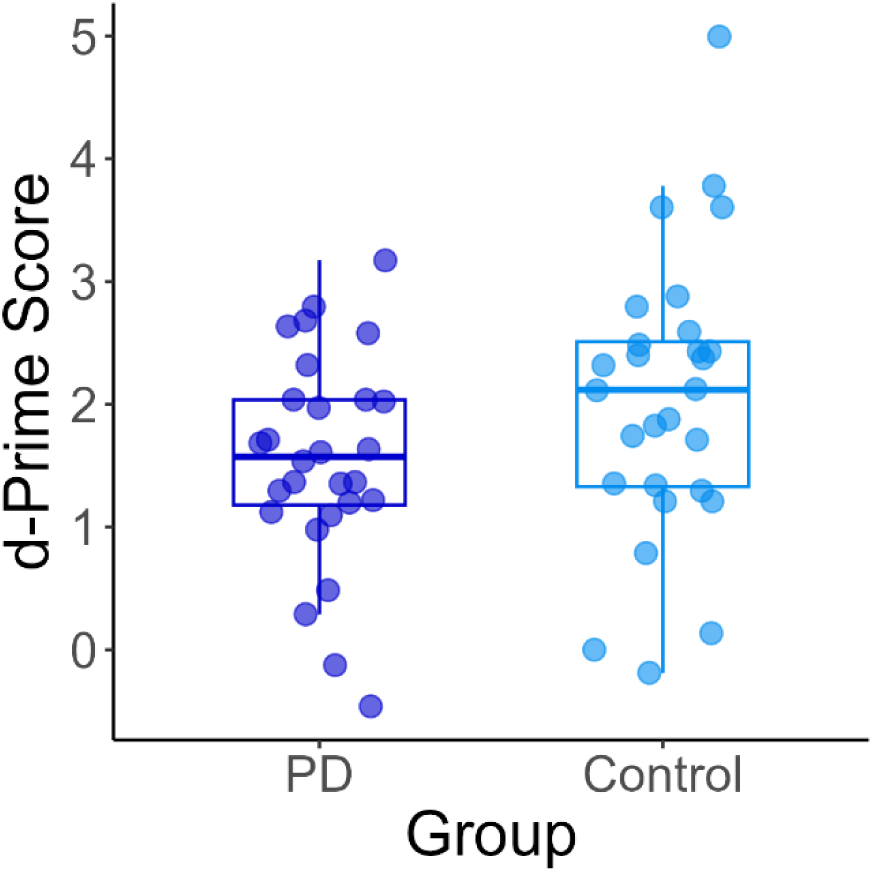
*d’* for the intensity discrimination task by participant group (participants with Parkinson’s disease [PD] in dark blue on left, control in light blue on right). The box plot shows median, 25^th^ and 75^th^ percentile scores, and values within 1.5 times the interquartile range (IQR) above and below the first and third quartiles. Each circle represents data from one participant.

#### Rate Discrimination

Figure 6 shows rate discrimination performance by stimulus type and participant group. A significant effect of stimulus type was found. Relative to normal speech, discrimination of time-reversed speech was significantly lower (*β* = -0.54, *t*(159.9) = -3.311, *p* = 0.001). d’ for non-linguistic tones (*β* = -0.30, *t*(159.9) = -1.837, *p* = 0.068) and the visual stimuli (*β* = -0.23, *t*(159.9) = -1.406, *p* = 0.162) were also lower than that of normal speech, but these differences were not significant. A significant effect of participant group was also observed, with control participants exhibiting significantly higher d’ than PD participants on average (*β* = 0.49, *t*(122.3) = 2.117, *p* = 0.036). No significant interactions were found between stimulus type and participant group (*p* > 0.05 for all).

**Figure 6.**
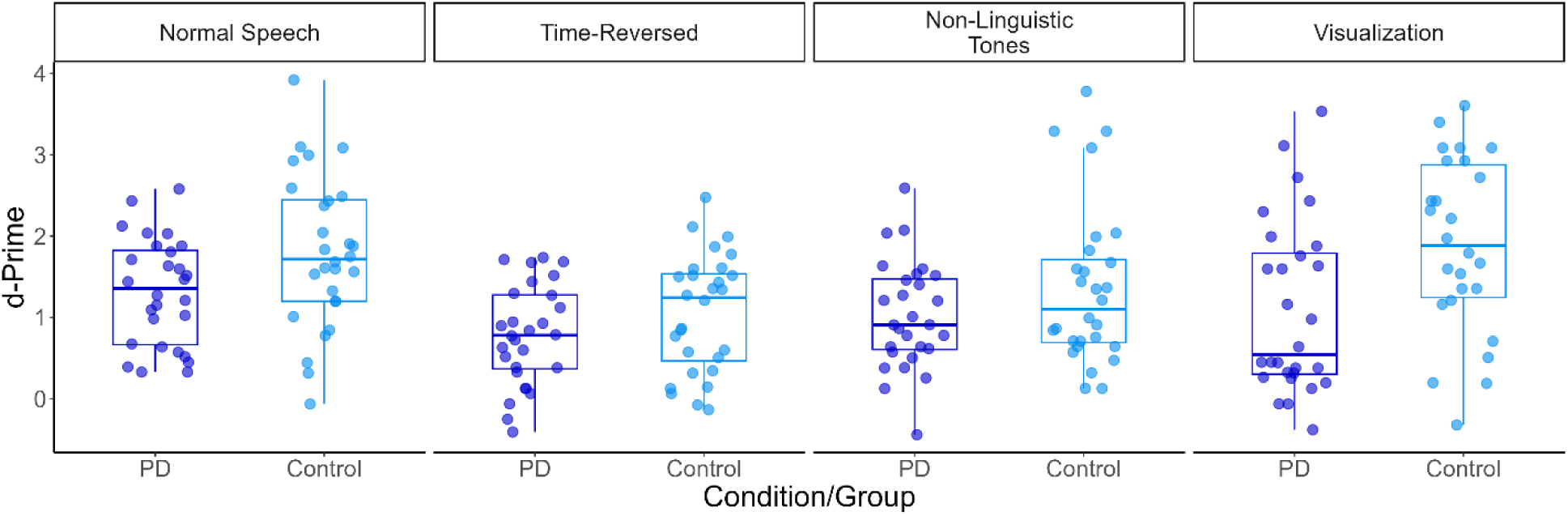
*d’* for the rate discrimination task by condition (paired box plots, labeled at top) and participant group (Parkinson’s disease [PD] in dark blue on left, control in light blue on right, within each condition). The box plot shows median, 25^th^ and 75^th^ percentile scores, and values within 1.5 times the interquartile range (IQR) above and below the first and third quartiles. Each circle is an individual participant.

### Disease Severity

Scores on the MDS-UPDRS indicated that the group of PD participants had overall mild symptom severity. Part I scores, indicating participant experiences with non-motor symptoms, ranged from 1 to 24 (m = 10.21, sd = 5.35). Part II scores, indicating participant experiences with motor symptoms, ranged from 1 to 25 (m = 10.17, sd = 6.87). Part III scores, experimenter assessment of motor function, ranged from 3 to 49 (m = 21.93, sd = 12.60). Despite the relatively mild severity of the PD participant group, individual differences among participants were found in both participant self-report (MDS-UPDRS Parts I+II) and experimenter examination (MDS-UPDRS Part III) of symptom severity.

Figure 7 shows the relationship between the *d’* values observed in the intensity discrimination task and the MDS-UPDRS scores obtained for PD participants. The relationship between the intensity *d’* value and Parts I+II of the MDS-UPDRS was not significant after accounting for hearing thresholds (*β* = -0.026, *p* = .089). However, there was a significant relationship between the intensity *d’* and Part III MDS-UPDRS, as illustrated in Figure 7, even after accounting for the effects of hearing thresholds (*β* = -0.031, *p* = .011).

**Figure 7.**
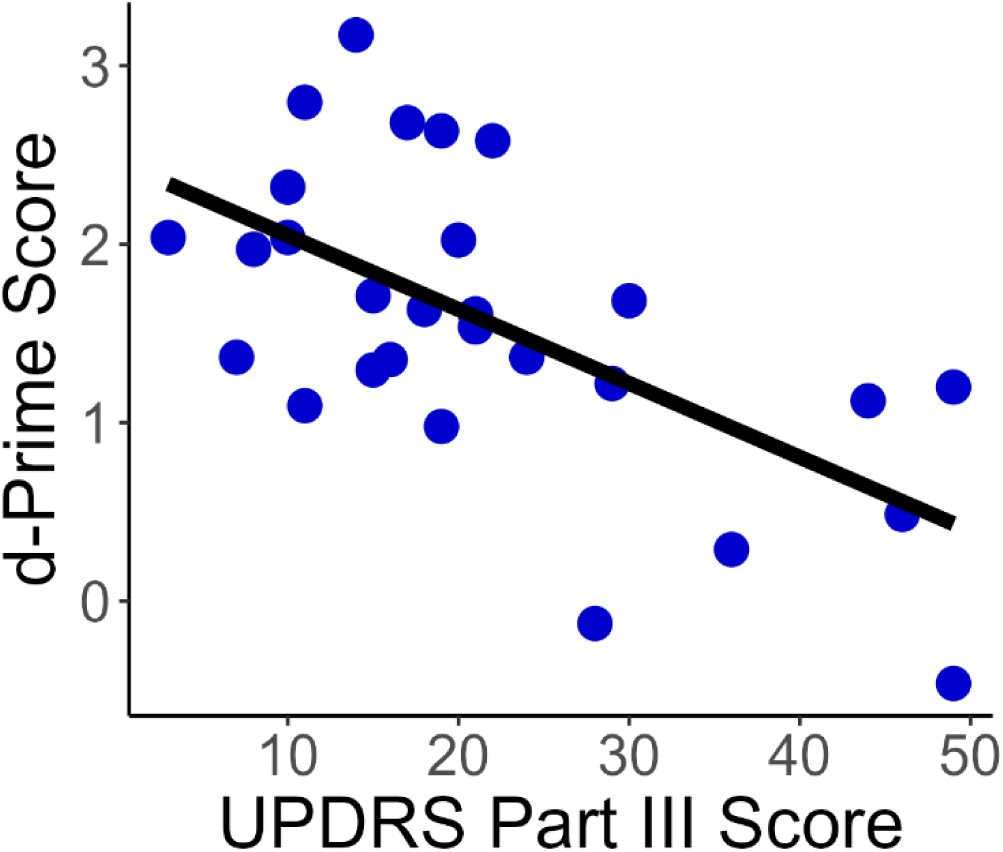
Scatterplot illustrating the relationship between performance on the intensity AXB discrimination task (as measured using *d’*) and the Part III sub score of the MDS-UPDRS test. Each dot is an individual participant with Parkinson’s disease (PD), while the black lines indicate linear best-fit lines.

For rate discrimination, however, linear modeling did not find a significant relationship between d’ and disease severity (Part I+II: *β* = -0.012, *p* = 0.363; Part III: *β* = -0.009, *p* = 0.401). Similarly, linear modeling revealed that MDS-UPDRS Part I+II and Part III scores were not significant predictors of accuracy on either the speech intensity (Part I+II: *β* = -0.003, *p* = 0.305; Part III: *β* < -0.001, *p* = 0.970) or the speech rate (Part I+II: *β* = -0.003, *p* = 0.780; Part III: *β* < -0.001, *p* = 0.887) learning tasks.

## Discussion

This study compared people with PD to age- and hearing-threshold matched controls in category learning of duration and intensity differences for speech stimuli, as well as discrimination of duration and intensity differences across a variety of modalities. We also examined relationships between PD severity and auditory perception. Although some participants had incomplete datasets, we included all usable data to maximize the sample size and increase generalizability of results. Relative to controls, people with PD showed significantly poorer performance in discriminating the duration, and therefore the perceived rate, of a variety of speech and non-speech stimuli. However, PD participants did not perform significantly differently from controls on the intensity discrimination task, nor in tasks of learning that employed rate and intensity manipulations. MDS-UPDRS Part III scores were also negatively correlated with performance on the intensity discrimination task, indicating that people with more severe PD had poorer performance on the intensity discrimination task for sentence stimuli.

### Category Learning of Speech Rate and Intensity

People with PD and age- and hearing-threshold matched controls did not differ in their accuracy of categorizing sounds that differed in intensity or duration. Performance for both groups improved over time, indicating that learning did indeed occur. Still, the groups did not differ in the speed of category learning. This finding was unexpected, given previous research that has implicated the basal ganglia in category learning, particularly of speech (see Lim et al., 2014 for review). Previous learning research in PD emphasizes procedural learning, as assessed with visual (e.g., Price & Shin, 2009) and motor (e.g., Gobel et al., 2013) sequence learning tasks. Several other investigations demonstrated that participants with PD performed more poorly on visual category learning relative to controls (e.g., Ashby et al., 2003; Shohamy et al., 2004). The only prior study of *auditory* category learning in PD to date showed that participants with PD performed more poorly than age-matched controls on tasks related to speech adaptation, but both groups performed similarly when learning to associate speech tokens of varying rates with colored squares – a category learning task (Heffner et al. 2022).

The present study expanded on the work by Heffner et al. (2022) by evaluating category learning at the sentence level in a new cohort of individuals with PD and matched controls. As in Heffner et al. (2022), we found similar results for PD participants and controls, despite a wealth of visual category learning literature that would predict differences for the two groups. The lack of learning differences between groups seem to be in line with a recent study conducted by Borrie et al. (2026), who found that individuals with PD and age-matched controls received the same benefit from perceptual training of dysarthric speech. Auditory learning, or at least specific types of auditory learning, may indeed be intact in PD. However, the current study excluded participants with mild cognitive impairment - whereas other investigations of category learning in PD did not. Previous observations of impaired category learning in participants with PD may have stemmed from the cognitive effects of the disorder, rather than directly from abnormal basal ganglia function. Indeed, Price & Shinn (2009) found that sequence learning in participants with PD was correlated with their cognitive flexibility, a measure of executive function. The current study’s exclusion of individuals with cognitive impairment may therefore explain the similarities in category learning between participant groups.

However, another potential explanation stems from the difficulty of the learning task. While on average both participant groups performed above chance (see Figure 4), individual participants in both groups performed at or below chance. The task, as it was constructed, did not (and could not) involve practice, as participants were expected to rely exclusively on feedback during the experiment in the form of a check mark indicating a correct response or an X indicating an incorrect response. The magnitude of rate or intensity differences between categories was also small – 10% time compression for rate learning and 2 dB for intensity learning. The learning task was specifically designed to be challenging so that performance did not approach ceiling too quickly. PD has been shown to affect feedback processing (Keitz et al., 2008), which made it a target for this paradigm. Still, if the task was too difficult in general, the control participants may not have benefited much from the feedback in a way to distinguish them from the PD participants.

### Discrimination of Speech Rate and Intensity

The finding of significant differences between participants with PD and controls for rate discrimination are consistent with previous studies suggesting people with PD exhibit challenges with timing discrimination (e.g., Su et al., 2022). Like previously published literature in the auditory domain (Forrest et al., 1998), participants with PD showed difficulty relative to age-matched controls in their ability to use linguistic information presented at different rates. Forrest et al. (1998), however, found that people with PD struggled to identify words played at a slower rate (but not at a normal or fast rate) relative to age-matched controls, whereas participants in the present study showed poorer ability to discriminate sentences played at many rates (both faster and slower than originally recorded). People with PD in the present study also performed more poorly than controls on duration discrimination for reversed-speech stimuli, non-linguistic tonal analogues, and even visual stimuli. Therefore, rate discrimination deficits were not limited to speech stimuli, in line with previous research with non-linguistic sounds (Grahn & Brett, 2009).

For intensity discrimination, although participants with PD demonstrated lower sensitivity relative to age- and hearing-matched controls, this group difference was not significant. This finding contrasts with results of Richardson and Sussman (2019). These divergent results are surprising because the previous study directly inspired aspects of the current methods: the step size for intensity in the current study was modeled on the point of greatest divergence between groups in the study performed by Richardson and Sussman (2019), which was 4 dB. However, speech stimuli differed between the two studies. Richardson and Sussman (2019) used single vowels, whereas the current study used whole sentences. The relatively more diffuse stimuli in the current study may at least partly explain why the overall discrimination performance was poorer than findings from Richardson and Sussman (2019). The d’ score for control participants was approximately 2 in the current study, whereas d’ averaged closer to 3 for Richardson and Sussman (2019). Differences in stimuli may have led to the divergence between studies.

Participants in the current investigation also differed in several ways from those in the study conducted by Richardson and Sussman (2019). First, participants in both groups of our study were on average older than participants in Richardson and Sussman. Average PTAs were similar between studies, but age-related changes in cognitive domains, such as attentional control or working memory, may have overridden perceptual ability and equalized performance between PD and control groups in our investigation. Second, and perhaps more likely, PD participants in the current study had generally more mild PD symptom severity than those in Richardson and Sussman (2019). Incorporating participants with greater PD severity in the current study may have resulted in a statistically significant difference in intensity discrimination between groups.

### Relationship to Disease Severity

PD symptom severity in the present study was measured using the MDS-UPDRS, a scale commonly used by neurologists to assess PD (Goetz et al., 2008). The Parts I+II score did not significantly predict participants’ learning or discrimination accuracy of rate or intensity. However, the Part III score significantly predicted intensity discrimination, even when statistically controlling for the effects of hearing threshold on intensity discrimination performance.

Interestingly, PD severity was not related to rate discrimination, despite PD participants demonstrating significantly lower accuracy across a range of rate-changed stimuli relative to control participants. It is possible that auditory rate and intensity deficits in PD are dissociable. Indeed, in PD speech production, changes in vocal intensity occur earlier on in disease progression than do changes in speech production rate (Logemann et al., 1978). In addition, the basal ganglia have been consistently linked to control and perception of timing (see Yin et al., 2014 for review) but not intensity. The network of neural coding required for stimulus intensity coding, and how it may be affected by PD, is less clear. Distinct physiological mechanisms therefore exist for rate and intensity coding for both motor and perceptual systems. Still, the PD participants in this study also demonstrated relatively mild symptoms, which may have limited observation of a relationship between disease severity and rate discrimination.

The finding that individuals with higher MDS-UPDRS Part III scores, but not Part I+II scores, had poorer intensity discrimination accuracy indicates a stronger relationship between auditory perception and gross motor function than between auditory perception and more general PD symptom experiences in everyday life. Although Part II of the MDS-UPDRS also assessed motor symptoms, it consists of self-report. Individuals’ perception of their symptoms differs from experimenter assessment of those symptoms. In addition, the current study used MDS-UPDRS scores as classification of disease severity, but information related to participants’ duration of PD or their levodopa equivalent daily dose was not collected nor compared to rate and intensity perception. Subsequent investigations could incorporate those variables in disease severity metrics to more precisely examine how PD symptoms, as well as dopamine-based medication dosage, relate to either intensity perception specifically or auditory perception more broadly. Such work will provide a better understanding of how dopamine networks may influence sensory perception.

As with all studies, there are limitations with the current investigation. First, the learning and discrimination tasks in this study utilized a single instance of a single sentence, perception of which is distinct from real-world processing of speech intensity and rate. The study results may therefore not accurately reflect participants’ everyday ability to perceive speech rate and intensity. For the learning tasks, participants were asked to categorize the acoustic information across an entire sentence; “category” learning may not be as robust for entire sentence-level stimuli as has been demonstrated for individual speech sounds (e.g., Heffner et al., 2019; Heffner & Myers, 2021). In addition, participants always performed the learning tasks first, and theoretically learned that certain ranges of intensity or duration were associated with a particular color square; this may have influenced discrimination, in that stimuli that had been presented within a category in the learning task may have been more difficult to discriminate relative to stimuli presented in different categories in the learning task. The perceptual tasks were also somewhat monotonous due to the small number of stimuli. Attention effects in both groups of participants may have influenced study results. Lastly, the PD participants in this study had relatively mild symptom severity and those with mild cognitive impairment were excluded, possibly affecting generalization of the study results to people with more severe PD symptoms.

Nevertheless, the results of this investigation provide additional insight into perception of acoustic speech features in PD. Future longitudinal studies could help to elucidate how auditory perceptual learning may change with disease progression. Subsequent investigations could also probe the extent to which speech perception challenges are connected to speech production in PD, as speech perception and production are fundamentally linked. For example, people who make clearer distinctions between vowel categories in their perception also make clearer distinctions in their production (Franken et al., 2017; Perkell et al., 2004). Based on the results of the current study, if individuals with PD have difficulty distinguishing temporal differences in their own speech production, this could explain some of the speaking rate changes observed with PD (see Blanchet & Snyder, 2009 for review). People with PD may simply not be able to detect differences between the rate that they intend to produce with the rate that is actually produced. An additional direction to pursue, then, may be perception of autophonic speech rate. If the perceptual deficits observed here are shared with perception of autophonic speech, this could strengthen the potential tie between speech perception and production networks – both in PD and more generally.

## Acknowledgements

Funding for this study was provided by an American Speech-Language-Hearing Foundation New Investigator Research Grant (NIRG) awarded to CCH, a University at Buffalo Blue Sky grant awarded to KT, CCH, and Eduardo Mercado III and startup funds provided to CCH by the University at Buffalo. The research was performed in the Cognition, Speech Perception, and Auditory Neuroscience Lab at UB, and we are grateful to lab personnel (including Raghad Azzam, Abbie Makofske, Abby Metzger, Tatianna Beutel, Alessia Mangoni, Alyssa DeCaro, Natalie Merriman, Madison Witt, Noora Somersalo, Tabitha Golda, Rylee Manning, and Rachel Maloney) for performing the research and to Jenna Crowell, Ivy-Jane Dimaculangan, Molly Krygowski, Nancy Stecker, Molly Stolze, and Celine Wan for support with hearing screenings. We would also like to thank Eduardo Mercado for assistance with project design, and to Kristina Milvae and Wei Sun for providing technical expertise.

## Data Availability Statement

The datasets generated during and/or analyzed during the current study are available in the Open Science Framework, available through this link: https://osf.io/am34k/?view_only=82d56c507abd46b991d94349d793ea54

